# Effect of light and darkness on the growth and development of downy mildew pathogen *Hyaloperonospora arabidopsidis*

**DOI:** 10.1101/2020.01.07.897215

**Authors:** Osman Telli, Catherine Jimenez-Quiros, John M. McDowell, Mahmut Tör

## Abstract

Disease development in plants requires a susceptible host, a virulent pathogen, and a favourable environment. Oomycete pathogens cause many important diseases and have evolved sophisticated molecular mechanisms to manipulate their hosts. Day length has been shown to impact plant-oomycete interactions but a need exists for a tractable reference system to understand the mechanistic interplay between light regulation, oomycete pathogen virulence, and plant host immunity. Here we present data demonstrating that light is a critical factor in the interaction between *Arabidopsis thaliana* and its naturally occurring downy mildew pathogen *Hyaloperonospora arabidopsidis* (*Hpa*). We investigated the role of light on spore germination, mycelium development, sporulation and oospore formation of *Hpa*, along with defence responses in the host. We observed abundant *Hpa* sporulation on compatible Arabidopsis under day lengths ranging from 10 to 14 hours. Contrastingly, exposure to constant light or constant dark suppressed sporulation. Exposure to constant dark suppressed spore germination, mycelial development and oospore formation. Interestingly, exposure to constant light stimulated spore germination, mycelial development and oospore formation. A biomarker of plant immune system activation was induced under both constant light and constant dark. Altogether, these findings demonstrate that *Hpa* has the molecular mechanisms to perceive and respond to light and that both the host and pathogen responses are influenced by the light regime. Therefore, this pathosystem can be used for investigations to understand the molecular mechanisms through which oomycete pathogens like *Hpa* perceive and integrate light signals, and how light influences pathogen virulence and host immunity during their interactions.

## Introduction

Environmental factors such as light, temperature and humidity play a significant role in the infection of plants by microbial pathogens and during disease development (Cheng et al., 2019). At the molecular level, adaptation to the environmental fluctuations is influenced by circadian timing mechanisms that undergo daily adjustment and act as a seasonal timer for diverse organisms, including plants and plant-associated microbes (Johnson et al., 2003). Light is the one of most significant environmental signals for circadian regulation (Dunlap et al., 2004). Many organisms have circadian regulation networks that operate through similar mechanisms. For plants, light is perceived by photoreceptors and act as a signal to regulate circadian genes (Millar, 2004; Franklin et al., 2005). Discrete light at different times of the day have been reported to have defined and particular effects on phase changes (Johnson et al., 2003). The circadian clock has also been shown to have a major effect on regulation of plant immunity (Karapetyan and Dong, 2018; Lu et al., 2017)

Light is known to have an effect on sporulation of several fungal and oomycete species, and the circadian clock of one fungal phytopathogen has been linked to the pathogen’s virulence programme (Hevia et al., 2015). Contrastingly, there are limited number of publications on the relation of light with development or virulence in oomycetes (Rumbolz et al., 2002). Early studies reported positive phototaxis of *Phytophthora cambivora* zoospores (Carlile, 1970) and the effect of humidity and light on discharge of sporangia of different oomycete pathogens (Fried and Stuteville, 1977; Leach et al., 1982; Su et al., 2000). Similarly, in *Plasmopara viticola,* the downy mildew pathogen of grapevine, continuous light did not have any effect on the growth of the mycelium and formation of sporangiophores, but the shape of sporangia was observed to be immature (Rumbolz et al., 2002). In the lettuce downy mildew pathogen *Bremia lactucae*, exposure to dark induced sporulation while light inhibited sporulation in a temperature-dependent manner: At low temperature, light was suppressive, however, with increasing temperature, the effect of suppression was decreased (Nordskog et al., 2007). Light was also suppressive of sporulation in *Peronospora belbahrii*, downy mildew of sweet basil, but light-dependent suppression of sporulation was enhanced at higher temperature. Light is also known to regulate the balance between asexual and sexual spore formation in *Phytophthora infestans,* causative agent of potato blight (Xiang and Judelson, 2014), in which exposure to constant light suppressed sporulation on plants and artificial media (Harnish, 1965). The mechanistic basis of light effects on oomycete virulence are largely unknown and likely to comprise a combination of light-regulated programmes for the host as well as the pathogen. It is also conceivable that the interacting organisms could directly influence each other’s circadian programs.

Oomycetes cause many important diseases of crops and in natural ecosystems (Kamoun et al., 2015). Much recent progress has been made in understanding plant-oomycete interactions through the development of reference plant-oomycete pathosystems that are amenable to genomic, genetic, and molecular approaches (Herlihy et al., 2019). One such pathosystem is comprised of the downy mildew pathogen *Hyaloperonospora arabidopsidis (Hpa)* and its natural host *Arabidopsis thaliana* (Coates and Beynon, 2010). Like many oomycetes, *Hpa* establishes an intimate relationship with its host by forming structures called haustoria, which are used to obtain nutrients from the plant. The *Hpa* life cycle is completed by the formation of aerial sporangiophores, which produce asexual spores, and by sexual oospores that are formed in infected leaves (Koch and Slusarenko, 1990). Because *Hpa* is an obligate biotroph, it requires its host to remain alive in order to complete its life cycle (Coates and Beynon, 2010). *Hpa* also redirects the host’s metabolism and suppress the host defence mechanisms (Herlihy et al., 2019). In *Hpa-Arabidopsis* interactions, it has been established that 16°C is the best temperature for *Hpa* sporulation under laboratory conditions (Dangl et al., 1992). However, the effect of different light/dark regimes on the sporulation of *Hpa* and the most productive light/dark time period for *Hpa* growth have not been reported. Elucidating the effect of light on the sporulation and growth of *Hpa* may also give some clue on whether there is a circadian regulation of its life cycle. Here, we report the effect of different light/dark regimes on the germination, mycelial development and sporulation of *Hpa*.

## Results

### Optimal light regime for *Hpa* sporulation

We began by testing how *Hpa* sporulation is affected by three different light (L) /dark (D) periods, representing day lengths commonly encountered by the plant and pathogen in natural environments. We used a compatible interaction between the *Hpa* isolate Emoy2 and a mutant in the Arabidopsis accession Columbia (Col) that inactivates the disease resistance gene *RPP4* (*Recognition of Peronospora parasitica gene 4*, Roux et al., 2011). Sporulation was quantified at four- and seven-days post-inoculation (dpi) under the following light regimes: 14h L / 10h D, 12h L / 12h D and 10h L / 14h D. Plants grown under all three regimes supported abundant sporulation, which increased between four and seven dpi (Figure 1). We observed only small, statistically insignificant differences in sporulation between the three regimes. We selected 12h L / 12h D as the reference time period for subsequent experiments.

**Figure 1.**
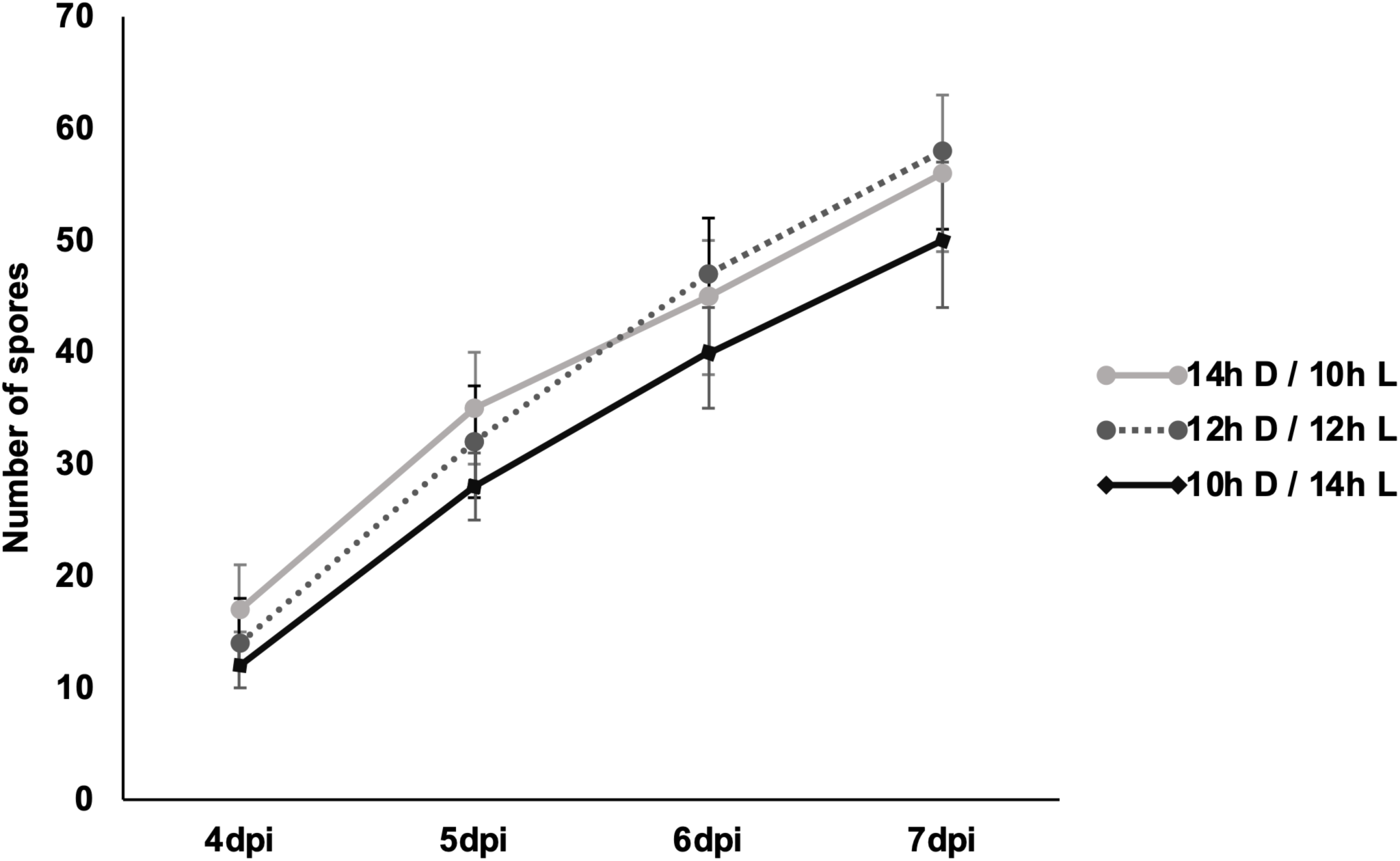
Optimization of the Light/Dark period for sporulation of *Hpa*. Three different light/dark periods were tested to compare the amount of *Hpa* sporulation sporulation. These periods were 14h L / 10h D, 12h L / 12h D, and 10h L / 14h D. Spores were harvested 4-7dpi and counted using a haemocytometer. Average and standard error of 3 replicates are shown. This experiment was repeated three times with similar results

### Exposure to constant light or dark suppresses sporulation of *Hpa*

The next set of experiments were designed to test how *Hpa* sporulation was affected by constant light or constant darkness. Four different light/dark conditions were compared to the 12h L / 12 h D reference: (1) Constant light exposure beginning after 3dpi; (2) constant dark exposure after 3dpi; (3) constant light exposure beginning immediately after inoculation; and (4) constant dark exposure beginning immediately after inoculation. As expected, abundant sporulation was observed between four and seven dpi on plants grown under the 12h L /12h D light regime (Figure 2A). Contrastingly, sporulation was dramatically reduced on seedlings exposed to constant light or dark after 3dpi. Moreover, sporulation was almost totally suppressed on plants grown under 7d constant light or 7d constant dark regime that commenced immediately after inoculation (Figure 2B). When infected seedlings were exposed to constant light or dark after 3dpi, there were hardly any new conidiophores and the amount of sporulation after 7dpi was the same as at 3dpi (Figure 2A). These experiments demonstrate that disruption of a normal light / dark regime can significantly affect the pathogen’s capacity to complete the asexual phase of its life cycle.

**Figure 2.**
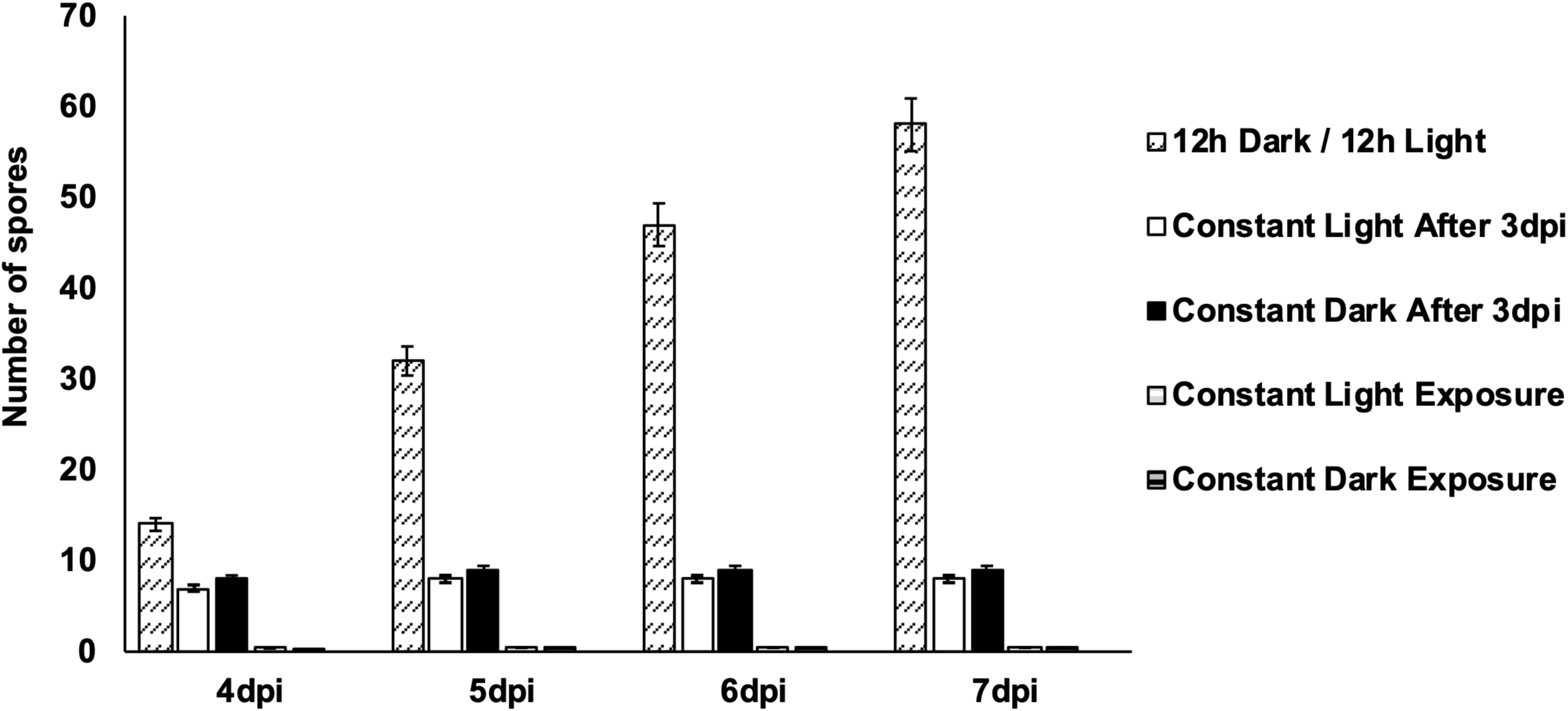
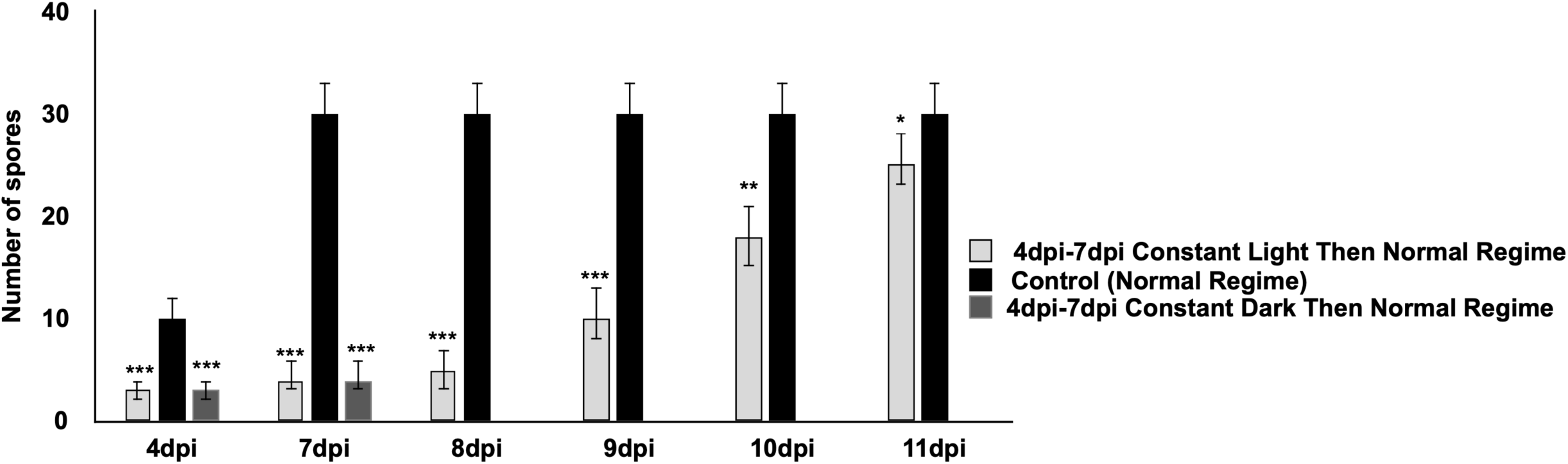
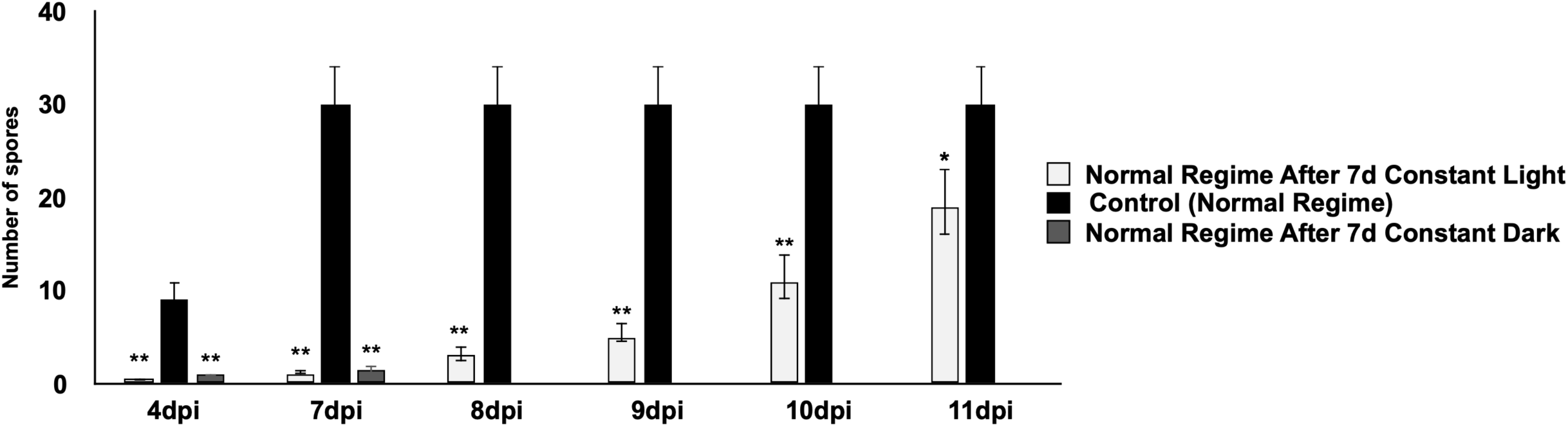
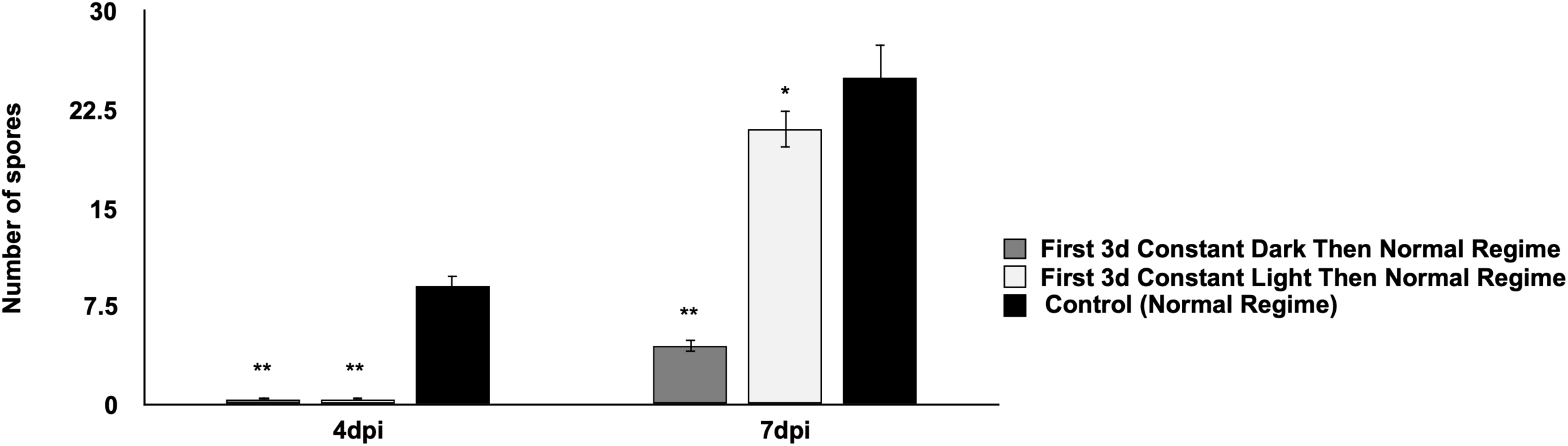
Amount of sporulation under different light regimes. **a)** Five different light/dark conditions were tested. These were 12h L / 12 h D (Dotted column), constant light after 3dpi (blank column), constant dark after 3dpi (black column), constant light exposure during 7d (grey/white column), constant dark exposure during 7d (black/grey column). Spores were harvested 4-7 dpi and counted using a haemocytometer. **b)** Samples were exposed to the reference light regime during the first 3 days (D/L black column), then were exposed to constant light (light grey column) or constant dark (dark grey column) over the subsequent 4 days (4dpi-7dpi). After end of the 7dpi, the samples were transferred to the reference light regime again and sporulation was recorded until 11dpi. **c)** Samples were exposed to constant light or constant dark for 7days immediately after inoculation. After 7dpi samples were transferred to the normal light regime again and sporulation was recorded until 11 dpi. **d)** Samples were exposed to constant light or constant dark beginning immediately after inoculation for 3 days, then shifted to a normal light regime, with sporulation recorded at 4 and 7 dpi. All experiments were repeated 3 times. All results were evaluated and compared statistically. *P < 0.05, **P < 0.01, ***P < 0.001, paired Student’s *t*-test

### Recovery from suppression of sporulation by constant light

We tested whether asexual sporulation could be restored by returning plants to the 12h light/ 12h dark after treatment with constant light or dark as described above. Interestingly, seedlings that were returned to a normal 12h L / 12 D regime after exposure to seven days constant light supported light sporulation 2 days after the shift and moderate sporulation after 4 days (Figure 2C). A similar recovery was observed in seedlings returned to the reference regime after treatment with constant light from 4-7 dpi (Figure 2C). Contrastingly, seedlings exposed to constant dark immediately after inoculation began to show a chlorotic phenotype after four days and the seedlings did not recover after shifting to normal light regime and no sporulation could be recorded (Figure 2C). Similarly, seedlings that were exposed to constant dark between 4dpi and 7dpi did not survive after 7dpi and thus no sporulation could be recorded (Figure 2B). When seedlings were exposed to constant light or constant dark beginning immediately after inoculation for 3 days, then shifted to a normal light regime, light sporulation was recovered 7dpi in samples exposed to constant dark. Abundant sporulation was observed 7dpi in samples exposed to constant light, similar to plants grown under a normal light regime (Figure 2D). These experiments demonstrated that the suppression of sporulation by constant light treatment of varying durations was not a permanent effect and that sporulation could be recovered by returning the plants to a normal regime.

### Different light conditions affect mycelial growth of *Hpa in* leaves

Considering that plants grown under constant light for 7d supported abundant *Hpa* sporulation after they were returned to a normal 12h L / 12 D regime (Figure 2), it seemed likely that mycelium may have grown inside the leaf during exposure to constant light but did not produce sporangia until a normal light regime was restored. To check this possibility, infected *At* seedlings were stained with trypan blue 3dpi. Trypan blue staining highlights mycelial growth along with sexual spore (oospores) that are produced in the interior of the leaf and asexual fruiting bodies (sporangia) that form on the exterior of the leaf.

In plants grown under the normal light cycle, mycelia had grown throughout cotyledons, sporangia had formed, and sporulation was observed over the whole surface of the cotyledon (Figure 3a). In contrast to the normal light cycle, in cotyledons exposed to constant light, there were extensive mycelia 3dpi and abundant oospores but no conidiophores (Figure 3b). These results indicate that vegetative growth and sexual sporulation can proceed under constant light, but asexual sporulation is suppressed.

**Figure 3.**
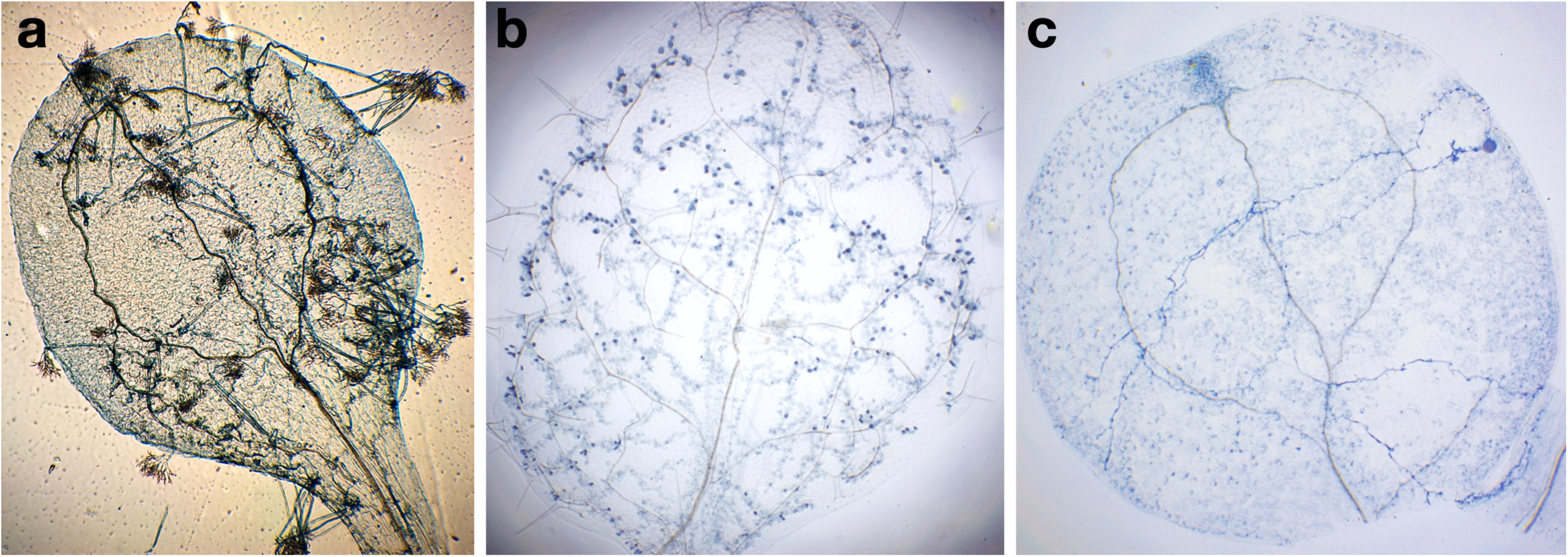
*Hpa* mycelial development in *At* leaves under different light regimes. **a)** Infected plants grown under the reference 12h light/ 12h dark cycle, **b)** Infected plants grown under a constant light regime, and **c)** Infected plants grown under constant dark regime. Infected *At* seedlings were stained with trypan blue 3dpi.

In cotyledons exposed to constant dark, less mycelial development was observed in those that were exposed to either a normal light cycle or constant light (Figure 3c). A small number of oospores were observed, similar to that observed under the constant light experiment.

To precisely assess *Hpa* growth *in planta*, we used a quantitative PCR assay in which *Hpa* DNA is quantified as a proxy for pathogen biomass. During evaluation over three days, mycelium biomass showed an increase in all groups (Figure 4). However, the lowest biomass was observed with constant dark exposure, whilst the constant light gave the highest biomass production in every day. Constant light conditions produced a significant increase in biomass compared to that observed with normal light conditions, especially at 3dpi. On the other hand, under constant dark conditions, biomass was significantly decreased compared to that obtained with the normal light conditions (Figure 4). Altogether, these results confirm that light is an important factor for vegetative growth and reproduction for *Hpa*.

**Figure 4.**
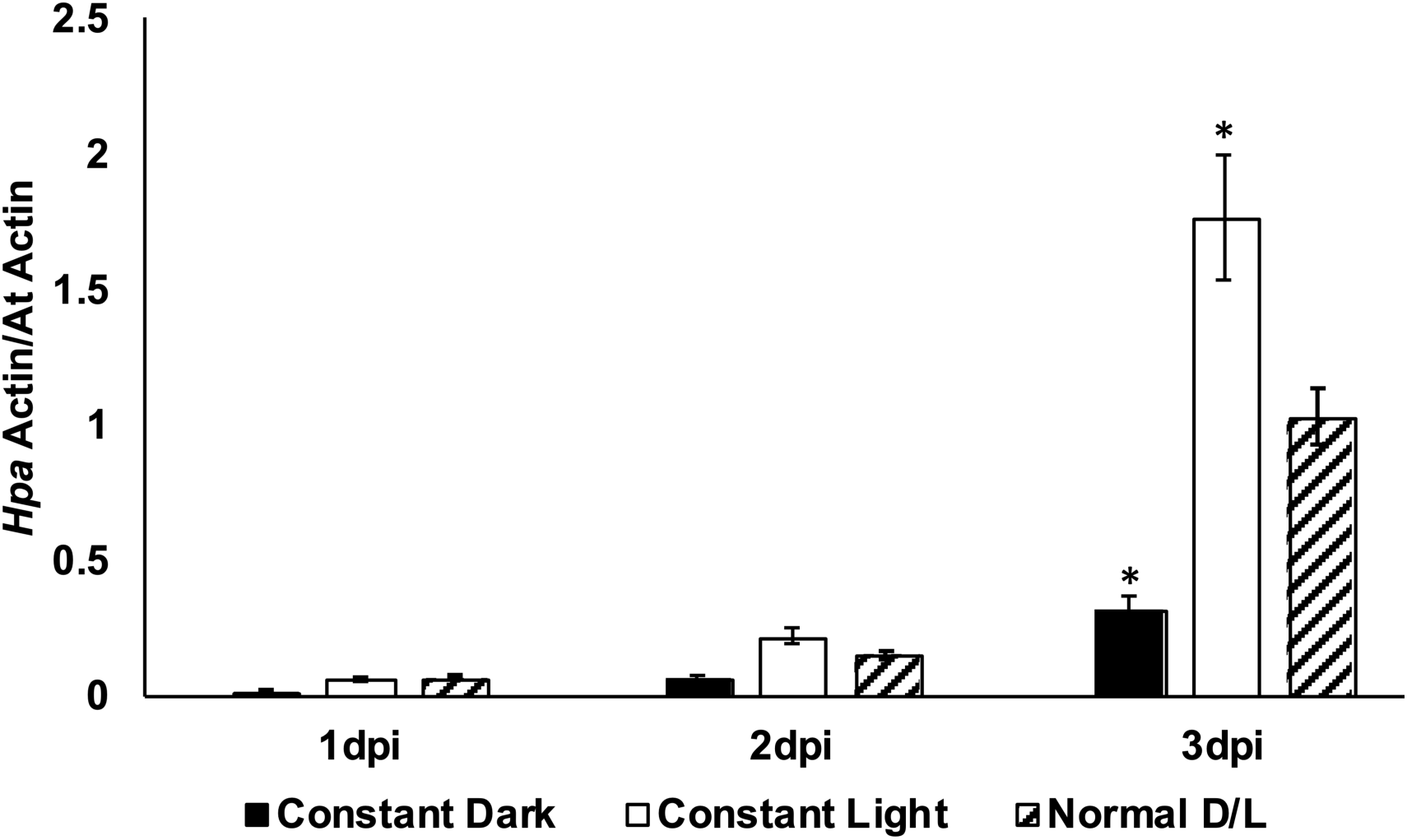
*Hpa* mycelial biomass production under different light regimes. Normal light (lined column), constant light (blank column) and constant dark (black column) regimes were applied. After *Hpa* inoculation, samples were taken every day from infected *At* leaves until 3dpi as mycelial growing phase is usually completed within first 3 days. In all samples, mycelial growth is calculated by qPCR and compared with each other. Student’s *t* test, *P < 0.05.

### Different light conditions affect spore germination

Because the light and dark affect *Hpa* vegetative growth and sporulation, we questioned whether the light or dark affect germination of spores and whether it is necessary to have a regular light/dark regime for germination. It is challenging to accurately quantify germination on plant leaves, because trypan blue staining and clearing during the early stages of infection eliminate spores on the leaf surface. Thus, cellophane strips were used for germination assays instead of seedlings.

The germination assay was first carried out with the reference light regime (12h L / 12h D). Under this regime, spores germinate after six to eight hours and a germ tube emerges (Figure 5a). After 12 and 24h, germ tubes have extended on the surface of the cellophane (Figure 5b and c). After 48 hours, formation of mycelial branches was obvious and most branches were laterally oriented as they covered the surface (Figure 5d).

**Figure 5.**
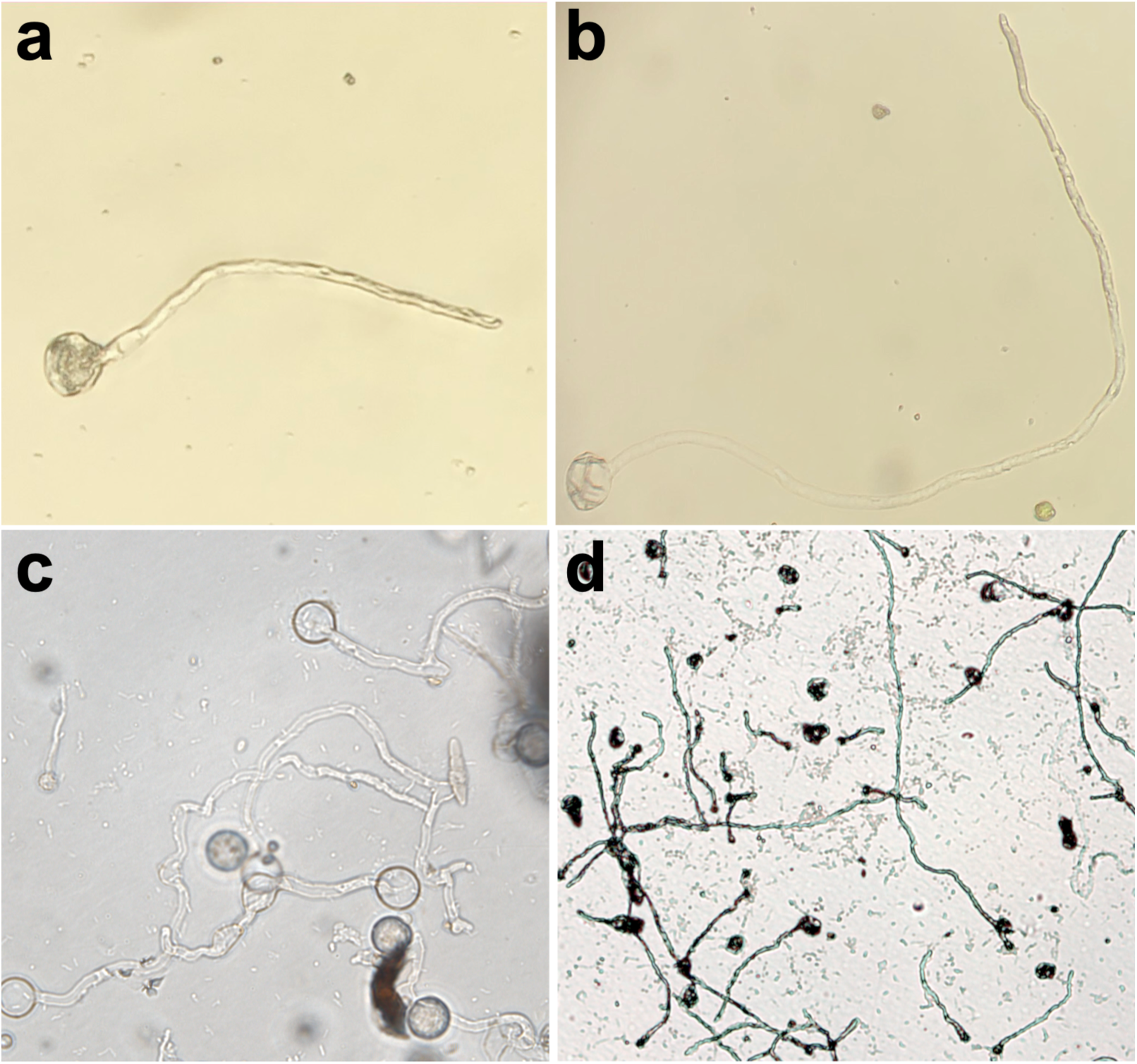
Germination of *Hpa* spores on cellophane under 12h L / 12h D regime. Spores were placed on cellophane strips and examined at regular intervals. **a)** after 6h, spore was germinated and germ tube was produced, **b and c)** after 12 and 24h, respectively, germ tube became longer, **d)** after 48h, lateral mycelial branches were obvious and hyphae began to cover the surface of the cellophane.

Germination using cellophane strips under constant light and constant dark was assessed in comparison to the reference light regime. The germination rate under the reference regime was 33% after 24 hours. The spores which were exposed to 24 hours constant dark showed a 22% germination rate, which was the lowest percentage observed within this time period. Under constant light after 24 hours, 37% of *Hpa* spores were germinated on cellophane (Figure 6). After 48h, the germination percentage increased for all treatments. The germination rate under constant dark was the lowest with 31%, the reference regime was 57% and constant light was 49%. After 72h, interestingly, the percentage of germination under constant dark and constant light was the same. However, in the reference light regime, germination increased and reached the highest percentage. At the end of 3d, germination seemed to be completed and spores appeared to have lost their viability. These results indicate that light is an important factor for spore germination independently of the host, and that optimal germination of spores occurs under a normal light/dark regime.

**Figure 6.**
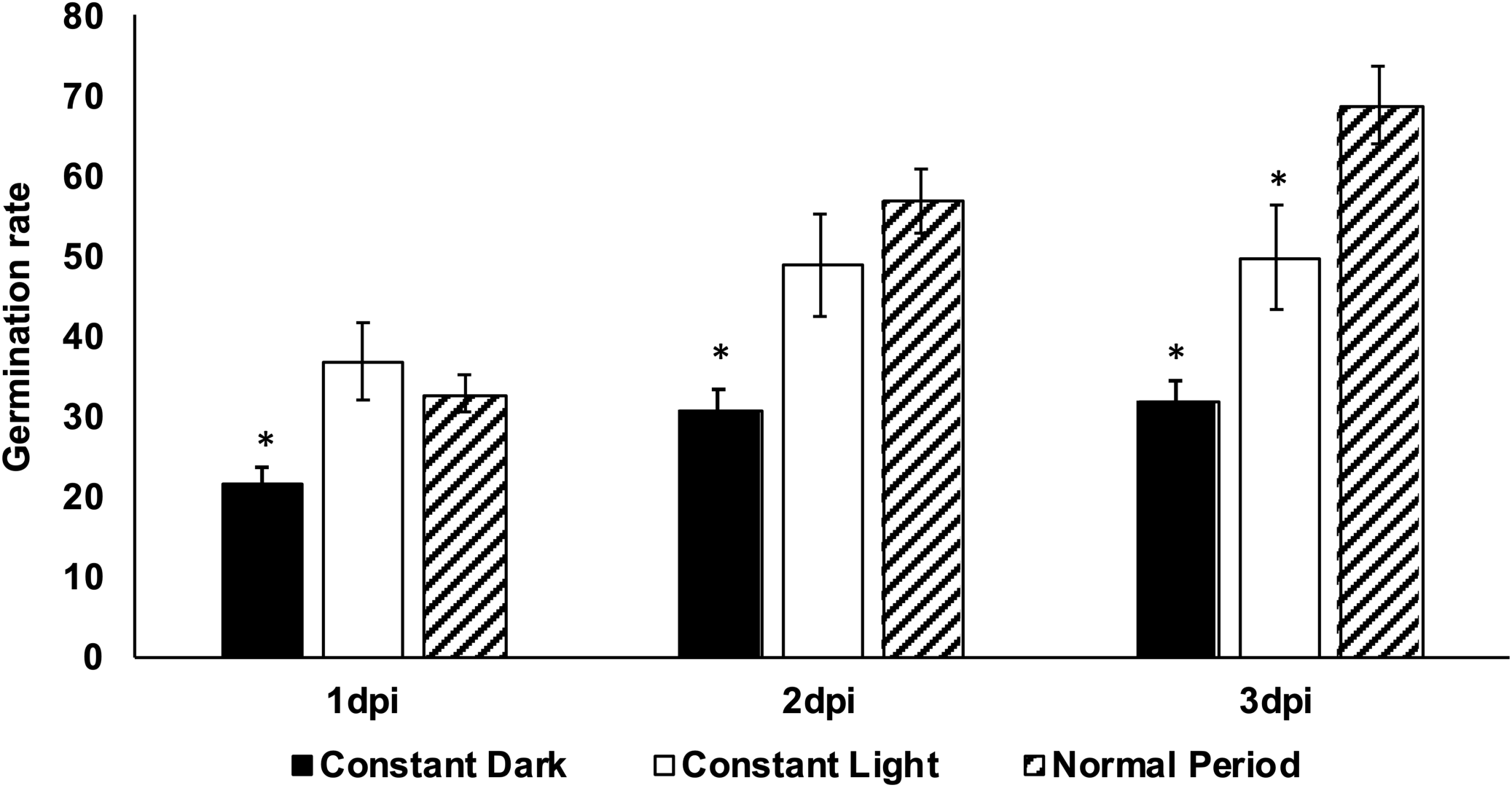
Germination rate of *Hpa* spores under different light conditions on cellophane. The spore germination on cellophane, which exposed to constant light (blank column), constant dark (black column) and 12h L/ 12h D (lined column) regime was determined after 24h, 48h and 72h. Values represent means of three experiments, and error bars correspond to the standard error of the means. Asterisks indicate statistically significant differences to the reference regime in two-tailed Student’s *t*-test (p <0,05).

### *Hpa* mycelial biomass growth is affected by inoculation time

If there is a synchronized circadian regulation of *Hpa* development and host defence, the inoculation time should be important for optimal colonization. Accordingly, previous reports have demonstrated that the time of day for inoculation can impact the degree to which *Hpa* can successfully colonize *Arabidopsis*, due at least in part to circadian upregulation of host immune responses during a time period that encompasses subjective dawn. Due to these observations, the optimal infection time for *Hpa* development was not obvious. Therefore, biomass productions between two inoculation times was compared using qPCR. Two zeitgeber time points were determined to observe the effect of day and night (or light and dark) on the development of pathogenicity. ZT0 refers to the beginning of daylight in an entrained cycle and ZT12 is the beginning of night, under experimental conditions of 12h L/ 12h D). One sample was inoculated at dawn (ZT0); beginning of the light period then followed by the dark period, therefore this sample was called L/D. The other sample was set up as the opposite; with inoculation at dusk (ZT12), called D/L. To determine the dynamic range of qPCR assays, we used an infection time course of virulent *Hpa*-Emoy2 on Col-*rpp4* (Figure 7). All samples showed a greater biomass of mycelium on the D/L cycle than L/D cycle and showed a regular increase in mycelium growth over the three days (Figure 7). The highest biomass was observed in the D/L cycle, where by three dpi, pathogen biomass in samples under the D/L cycle was approximately 56% higher than in samples inoculated under L/D cycle. The results suggest that initiating infection at dusk promotes a higher degree of virulence than initiating infection at dawn.

**Figure 7.**
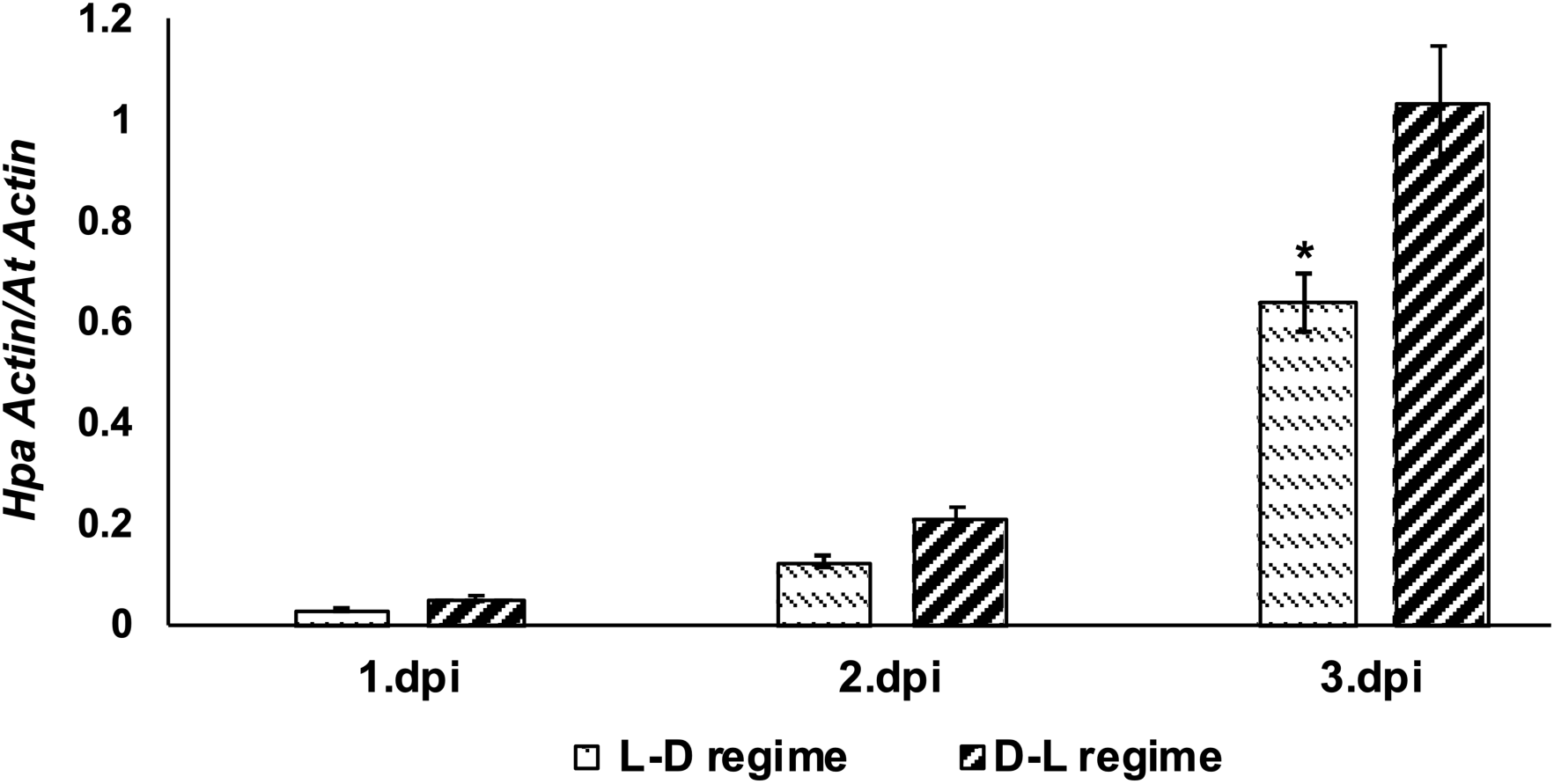
Effect of inoculation time on *Hpa*-Emoy2 biomass on Col-*rpp4*. Col-*rpp4* seedlings were infected with *Hpa-*Emoy2 at dawn (ZT=0), labelled LD (white dotted column) or at beginning of the dark period (ZT=12) labelled DL (black lined column). At the end of each day, samples were taken and biomass was calculated using qPCR and compared with each other. Student’s *t* test, *P < 0.05.

### Continuous light/dark regimes activate an immune response biomarker in plants

Plant immune responses are characterized by the activation of a set of pathogenesis-related (*PR*) genes (Ward et al., 1991; Uknes et al., 1992). A *PR1promoter-GUS* reporter gene is considered to be a valid marker gene for activation of immunity in *A. thaliana* (Uknes et al., 1992), enabling transgenic *Arabidopsis PR1-GUS* plants to be employed to detect activation of immune responses. We set up an experiment to test whether the reporter gene activity can be induced by constant light and constant dark treatment with the transgenic plants containing *PR1-GUS.* In this assay, when the *GUS* reporter gene is activated by any stress factor, the plant tissues are observed to be stained blue.

In *Arabidopsis* seedlings grown under normal light regime with no pathogen infection, there were no blue stained cells indicating that the *GUS* gene was not induced under this condition, as expected (Figure 4a). Contrastingly, seedlings exposed to constant light (Figure 4b) and constant dark (Figure 4c) for 72h showed *GUS* activity, indicating constant light and dark regimes trigger immunity. These results indicate that induction of immunity under constant light or dark exposure could contribute to the suppression of the sporulation of *Hpa*.

## Materials and methods

### Plant lines, pathogen isolates and propagation

*Hyaloperonospora arabidopsidis* isolate Emoy2 were maintained on Ws-*eds1* (Parker *et al*., 1996) or Col-*rpp4* (Roux et al., 2011). Maintenance and preparation of inoculum for experiments was performed as described previously (Tör *et al*., 2002; Woods-Tör 2018). Transgenic *PR1-GUS* lines were obtained from Xinnian Dong.

### Sporulation assay

Inoculated Col-*rpp4* seedlings were exposed to 3 different light (L) /dark (D) periods; 12h L/12h D, 14h L/10h D and 10h L/14h D for 7 d at 16°C and the amount of sporulation was assessed.

Another experiment was designed to understand the effect of extreme light regimes on *Hpa* sporulation. The inoculated samples were exposed to 4 different light regimes; 7 d constant light, 7 d constant dark, constant light after 3dpi and constant dark after 3dpi, and control light/dark regime (12h L/ 12h D) was also included. As a light source, white fluorescent bulbs (300 mmol m±2 s±1, 10 Osram HQIL 400 W-lamps plus four Osram L40/ W60 fluorescent bulbs; Osram, Berlin, Germany) were used. To quantify sporulation, 10 infected seedlings from each replicate were taken and placed into an Eppendorf tube containing 250µl H20. Samples were vortexed and conidiospores were counted using a haemocytometer. All experiments had minimum three replicas and were repeated 3 times. All results were evaluated and compared statistically.

### Trypan blue staining

Cotyledons of 7 d old *Col-rpp4* were spray inoculated with *Hpa*-Emoy2 and were exposed to a normal 12h L / 12h D cycle, constant light or constant dark and examined at 3 dpi after staining with Trypan Blue as described at below;

Seedlings were taken from infected samples at the 0 hrs, 12 hrs, 1d, 2d, 3d, 4d, 5d, 6d, 7d post inoculation (dpi). Infected leaf segments were placed in an Eppendorf tube, covered with 1 ml or enough amount trypan blue solution (10 g phenol, 10 ml glycerol, 10 ml lactic acid, 10ml water and 0.02 g of trypan blue (Merck) in ethanol (96%; 1:2 v/v) and boiled at 100 °C for 1 min. The leaf segments were then de-stained for an hour in chloral hydrate (2mg/ml) (Sigma). All steps were carried out in a fume hood. Pathogen structures were viewed under a CARL Zeiss Axioskop 4 plus microscope.

### GUS assay

Transgenic *PR1-GUS* lines were used. Three-week-old seedlings were exposed to constant light or dark for 1 to 3 days. Then, seedlings were transferred to 24 well replica plates that contained 1 ml X-Gluc histochemical staining solution (50 mM X-Gluc in 50 mM NaPO4 pH 7.0) and incubated overnight at 37 ^0^C. After staining, leaves were treated with 70% methanol up to 4 h. The samples were washed with ethanol, immersed in glycerol and tissues were examined for GUS staining under dissecting microscope.

### Germination assay using cellophane

The germination assay using cellophane on MS (Murashige and Skoog, 1962) was carried out as described (Bilir et al., 2019). Sterile pieces of cellophane were placed on the surface of MS agar in the flow cabinet. *Hpa* spore solution was prepared and centrifuged, all spores collected, and the pellet was then resuspended in sterile water. Approximately 10 µl spore solution were dropped on each piece of cellophane. Plates was grouped and put in the 3 different incubators; constant light, constant dark and 12h L/12h D regime at 16°C during 72h and examined every 12h under microscope. The number of germinated *Hpa* spores was counted using a haemocytometer.

### Determining biomass growth using qPCR

The biomass of mycelium produced by *Hpa*-Emoy2 up to 3dpi was measured from samples exposed to three different light regimes by Real-Time Quantitative PCR (RT-qPCR). The *Hpa-Actin* gene and *At-Actin* gene were used for quantification and its relative protocol was followed as described (Anderson and McDowell, 2015). After Col-*rpp4* seedlings were inoculated with *Hpa*-Emoy2, samples were separated and placed under normal (D/L), constant light and constant dark regime as three different groups. Every 24h, samples were taken, and their DNA extracted and calculated with qPCR as described (Livak and Schmittgen, 2001). Sequences of primers used were Hpa-Actin/F 5’-GTTTACTACCACGGCCGAGC-3’, Hpa-Actin/R 5-CGTACGGAAACGTTCATTGC-3’, At-Actin/F 5’-AGCATCTGGTCTGCGAGTTC-3’, and At-Actin/R 5’-ACGGATTTAATGACACAATGGC-3’.

### Statistical analysis

For statistical analysis, paired Student’s *t*-tests were performed on data obtained from plant infection and germination assays.

## Discussion

Using *Hpa-Arabidopsis* reference system, we showed that light regimes significantly affect several stages of the *Hpa* disease cycle, including spore germination, mycelial development, oospore formation and sporulation. We also obtained preliminary result suggesting that light regimes can also influence the immune status of the host. These observations complement recent studies showing that the plant circadian clock system regulates the immune system in the interactions between Arabidopsis and *Hpa* (Wang *et al*., 2011; Zhang *et al.,* 2013). However, the previous studies focused mainly on incompatible interactions with resistant plant hosts and did not address how light might impact *Hpa* in a disease-susceptible host. Therefore, this work was undertaken to investigate the effect of light on a virulent *Hpa* isolate.

Our first observation was that an entrained light/dark cycle was necessary for *Hpa* to efficiently complete its life cycle in the host. We observed only minor differences in spore production from plants grown in three different light regimes (14h L / 10h D, 12h L / 12h D and 10h L / 14h D; Figure 1) and selected 12h L /12h D as a reference regime for pathogen for ongoing experiments.

We then exposed plants to constant light or dark regimes, commencing after three dpi for four days total or immediately following infection for seven days. All of these regimes had a suppressive effect on sporulation (Figure 2). Similar inhibitory effects of light on sporulation of fungal and oomycete pathogens, including downy mildews, have been reported for decades [referenced in the Introduction and reviewed in (Rotem et al., 1978)]. However, these studies generally have not directly addressed whether constant light inhibited vegetative (mycelial) growth in planta and/or sporulation. We assessed *Hpa* growth in the leaves with quantitative PCR and with Trypan Blue staining. Both assays indicated that *Hpa* growth was moderately inhibited in dark grown plants but was not inhibited in plants exposed to constant light. Indeed, constant light supported higher levels of *Hpa* biomass than that in normal light/dark or constant dark regimes. In trypan blue staining experiments, in the normal light regime, the initial stages of conidiophore development were observed at 3 dpi, while in constant light experiments, abundant oospore formation was observed 3dpi. This may indicate that constant light exposure could inhibit asexual sporangiophore development while acting as an inducer of oospore formation.

Interestingly, this apparent inhibition of asexual sporulation by constant light or dark was reversible: plants that were returned to the reference light regime after four days of constant light or dark could support abundant sporangiophore production. Similar observations have been reported for other downy mildew pathogens, for which a “recovery” period of four hours in the dark was sufficient to enable sporulation (reviewed in Rotem et al., 1978). The mechanism behind this recovery is unknown but was postulated at the time to involve enzymatic degradation of a light-induced “antisporulant”. Such hypotheses can now be tested with the experimental tools of the *Hpa*-Arabidopsis pathosystem.

In this context, we tested whether constant light or dark-treatment was sufficient to activate the plant immune system in the absence of pathogen infection. Using transgenic plants containing a fusion of *PR1* promoter to a *GUS* reporter gene, it was clear that after 24h, the *PR1* promoter was activated by 24h constant light and 48h constant dark (Figure 4). These results are similar to those reported in previous publications (Evrard et al., 2009). It has been reported that plant defence responses and HR-associated programmed cell death triggered by pathogen is activated by light in tobacco (*Nicotiana tabacum*), rice (*Oryza sativa*), and *Arabidopsis*, and the activation of inducible resistance is dependent on phytochrome functions (Guo et al., 1993; Chandra-Shekara et al., 2006). The blue light receptor cryptochromes (*CRY*) and red/far-red light photoreceptor phytochromes (*PHY*) work together in *Arabidopsis* and they regulate many light-controlled defence responses and entrainment of the circadian clock. The photoreceptor gene *CRY1* regulate systemic acquired resistance (SAR) positively and in the *cry1* mutant, salicylic acid (SA)-induced pathogenesis-related gene *PR-1* expression is reduced but enhanced in *CRY1-ovx* (*CRY1-overexpressor*) plants under light conditions (Liang and Hong-Quan, 2010).

We also tested whether the timing of inoculation affected *Hpa*’s capacity to colonize the plant. A previous report demonstrated that effector-triggered immunity and basal immunity against *Hpa* is more efficient early in the day (Wang et al., 2011), and we confirmed this observation by using a different virulent isolate of *Hpa*. Our results demonstrate that plants inoculated at dusk supported significantly more mycelial growth than plants inoculated at dawn, even at three dpi. Our experiments do not point directly to an underlying mechanism, but we hypothesize that this might reflect a difference in timing of basal defense mechanisms that limit growth of virulent *Hpa*. Wang et al (2011) noted that SA-dependent gene expression was stronger in the day than at night; accordingly, it was reported that morning and midday inoculations lead to higher salicylic acid accumulation, quicker and more intense *PR* (pathogen-related) gene activation and expression, and HR responses than inoculations in the dusk or at night (Griebel and Zeier, 2008). These previous reports on different systems support our data and help to explain why night time inoculation is more efficient than day time inoculation (Figure 8).

**Figure 8.**
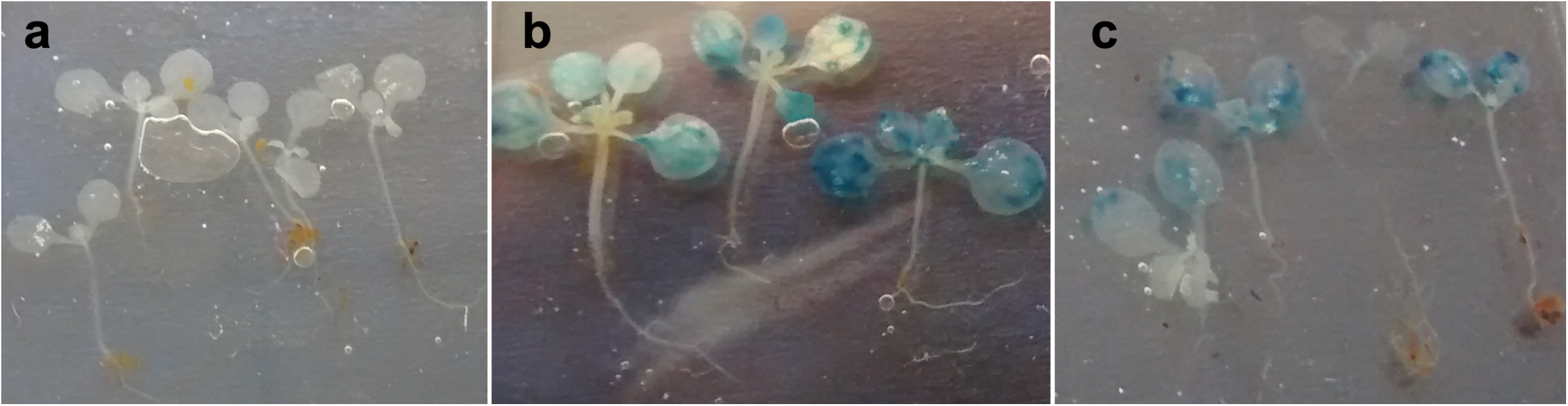
*GUS* expression in *At* seedlings exposed to different light regimes. **a)** Seedlings grown under normal 12h L / 12h D regime. **b)** Seedlings exposed to constant light. **c)** Seedlings exposed to constant dark. After 48 hours of exposure to these regimes, histochemical GUS assays were carried out. These experiments were repeated 3 times with similar results.

It is important to emphasize that all experiments involving *Hpa* grown *in planta* could reflect influence of light on both the pathogen and the host. Fungal and oomycete pathogens have been shown previously to incorporate light perception into their development and virulence programs. For example, 48h constant white light exposure inhibits sporulation of *P. infestans* on potato or agar plates (Xiang and Judelson, 2014). Because *Hpa* is an obligate pathogen that can only complete its life cycle on a compatible *Arabidopsis* host, we cannot directly assess how light influences sporulation apart from the host. However, our *in vitro* spore germination assay indicates that light does affect the *Hpa* life cycle and suggests that *Hpa* can perceive light.

In conclusion, we have reported several lines of evidence that light is a critical factor during development of downy mildew disease on *Arabidopsis* and can influence responses in the pathogen and the host. We can now exploit this system to understand the mechanistic basis of these effects, using the well-developed tools for *Arabidopsis* in combination with a new protocol for reverse genetics in *Hpa*. Our future studies will focus on circadian regulation on both the host and pathogen side. While it is well-established that circadian regulation of host immunity is an important factor in immunity against *Hpa* and other pathogens in *Arabidopsis*, the role of circadian regulation in oomycete virulence is unexplored and therefore could be an enlightening area for future inquiries.

## Author contributions

MT and OT planned and designed the research. OT and CJQ conducted the laboratory work. MT, OT and JMM analyzed and interpreted the data and wrote the manuscript.

## Data availability statement

The data that support the findings of this study are available from the corresponding author on reasonable request.

